# A Robust Method for Collecting X-ray Diffraction Data from Protein Crystals across Physiological Temperatures

**DOI:** 10.1101/2020.03.17.995852

**Authors:** Tzanko Doukov, Daniel Herschlag, Filip Yabukarski

## Abstract

Traditional X-ray diffraction data collected at cryo-temperatures have delivered invaluable insights into the three-dimensional structures of proteins, providing the backbone of structure-function studies. While cryo-cooling mitigates radiation damage, cryo-temperatures can alter protein conformational ensembles and solvent structure. Further, conformational ensembles underlie protein function and energetics, and recent advances in room-temperature X-ray crystallography have delivered conformational heterogeneity information that is directly related to biological function. The next challenge is to develop a robust and broadly applicable method to collect single-crystal X-ray diffraction data at and above room temperatures and was addressed herein. This approach provides complete diffraction datasets with total collection times as short as ~5 sec from single protein crystals, dramatically increasing the amount of data that can be collected within allocated synchrotron beam time. Its applicability was demonstrated by collecting 1.09-1.54 Å resolution data over a temperature range of 293–363 K for proteinase K, thaumatin, and lysozyme crystals. Our analyses indicate that the diffraction data is of high-quality and do not suffer from excessive dehydration or damage.

## Introduction

Structures obtained from X-ray diffraction of cryo-cooled protein crystals have arguably provided the most impactful contributions of physics to biology. It is now routine to visualize the fold, intramolecular interactions, and binding sites of proteins–information with profound implications for the understanding of protein structure, function, and evolution (e.g. Berg *et al*., 2002; Brändén & Tooze, 1999; Fersht, 2017; Ufimtsev & Levitt, 2019; Wlodawer *et al*., 2008); and the thousands of examples of protein structures, along with simplified energetic rules, have led to our current ability to predict structure from sequence for many proteins and to design proteins that form specified folds in many cases (Huang *et al*., 2016; Kuhlman & Bradley, 2019; Marks *et al*., 2012).

In contrast, our ability to predict the energetics of protein folding, binding, and function is limited. This contrast appears to arise from fundamental principles of physics: free energy, which specifies preferred states and their relative occupancy, is determined from relative energies of states within the protein conformational ensemble. Traditional X-ray crystallography provides structural information at 100 K, but temperatures below the so-called glass transition (generally in the 180-220 K range) can alter protein conformational heterogeneity, the experimental manifestation of conformational ensembles, and quell function (Fraser *et al*., 2009; Halle, 2004; Juers & Matthews, 2001; Keedy *et al*., 2014; Sandalova *et al*., 1999). Further, and likely of more general importance, traditional X-ray crystallography models provide limited conformational heterogeneity information (Ringe & Petsko, 1985; Petsko, 1996; Furnham *et al*., 2006). Underscoring the need for in-depth detailed information about conformational heterogeneity, there has been considerable discussion about the tuning of protein motions and conformational heterogeneity to suit physiological temperatures (Feller, 2010; Fields *et al*., 2015; Siddiqui & Cavicchioli, 2006; Elias *et al.*, 2014).

Unlocking the potential of X-ray crystallography to provide conformational heterogeneity information at physiological temperatures that can more directly be related to native conformational ensembles, energetics, and function, requires an ability to routinely obtain high-quality X-ray diffraction data at physiological temperatures. While historically X-ray diffraction data had exclusively been collected at room temperatures (RT), cryo-cooling crystals allowed important improvements in data quality, speed of data collection, and amount of information that could be obtained from a single crystal (Hope, 1988), and cryo-temperature data collection quickly overtook protein X-ray crystallography. Nevertheless recent advances in X-ray sources, optics, and detectors has led to a renaissance in RT X-ray crystallography data collection, and parallel methods development has enabled conformational heterogeneity information to be obtained from the RT diffraction data and to be related to function (Fraser *et al*., 2011, 2009; Keedy *et al*., 2014, 2018; van den Bedem *et al*., 2009; Lang *et al*., 2010).

Although technical and methodological progress in collecting higher temperature data, demonstrated that physiological temperature data collection is possible, these experiments remain challenging (Rajendran *et al*., 2011). An ability to routinely obtain these data is needed to expand the usage and thus impact of RT X-ray crystallography. In addition, the ability to obtain data across the range of physiological temperatures would allow models for the evolutionary tuning of protein function and the origins of protein conformational heterogeneity to be tested and new models to be developed. Ultimately, with sufficient data, these approaches, coupled with computational advances, will extend our abilities from predicting structures to predicting conformational ensembles, the latter being related to the energy of the system via the laws of statistical mechanics.

Here we present a broadly applicable and robust method for efficiently collecting single crystal X-ray diffraction data at and above room temperatures from crystals of ≥0.3 mm on each side at synchrotron beamlines. We present the technical aspect of the instrumentation and data collection strategy that have allowed us to obtain single crystal X-ray diffraction data at beamline 14-1 (BL 14-1) at the Stanford Synchrotron Light Source (SSRL) and can be generalized to other beamlines. With this approach, we can obtain complete diffraction datasets of high quality with total collection times as short as ~5 sec, dramatically increasing the amount of high-quality data that can be collected during allocated beam time at experimental X-ray crystallography stations.

## Experimental

### Obtaining crystals for X-ray diffraction at and above room temperature

*Tritirachium album* proteinase K (catalog # P2308), *Thaumatococcus daniellii* thaumatin (catalog # T7638), and hen egg lysozyme (catalog #L4919) were purchased from Sigma and crystallized at room temperature as previously described (https://www1.rigaku.com/en/products/protein/recipes) using a hanging drop (proteinase K and lysozyme) and sitting drop (thaumatin) setups.

Crystals are more sensitive to radiation damage at room temperatures than at cryo temperatures (see below), and the diffractive contribution from a unit cell is destroyed by a lower number of absorbed photons than at cryo temperatures (Garman & Weik, 2017; Garman & Owen, 2006; Nave & Garman, 2005; Roedig *et al*., 2016; Southworth-Davies *et al*., 2007; Warkentin *et al*., 2011; Warkentin & Thorne, 2010); thus, collecting X-ray diffraction data at and above room temperature to resolutions approaching resolutions from cryo-cooled crystals requires a larger number of unit cells (and a correspondingly larger crystal). Here we used crystals of dimensions ≥0.3 mm on each side (i.e. 0.3–0.4 mm); data collected from smaller crystals was neither complete nor to high resolution, with diffraction statistics indicating excessive radiation damage, as anticipated (not shown). When compared to micrometer-sized crystals needed for serial XFEL crystallography, and the millimeter-sized crystals required for neutron diffraction studies, the crystals used here were of ‘intermediate’ size and appeared optimal for data collection at and above room temperature. To maximize diffraction intensity while minimizing the number of absorbed photons per unit cell, we matched the beam and crystal size as closely as possible.

### Achieving high-temperature capabilities and temperature control

To enable high temperature data collection at the Stanford Synchrotron Radiation Lightsource (SSRL) BL 14-1, an Oxford Cryosystems Cryostream 800 model N2 heater/cooler with a temperature range of 80–400 K (temperature stability of 0.1 K) was installed. The nozzle was aligned coaxially to the sample holding pin (**Figure 1A**), and the temperature at the crystal position was confirmed with a type J thermocouple on Omega HH23 microprocessor thermometer. Because the physical properties of protein crystals deteriorate with high temperature exposure, we used crystal annealer in a ‘sample protective mode’ to control the crystal exposure to the N2 stream as follows: after the desired (high) temperature of the N2 stream is achieved and prior to mounting the sample in the N2 stream, the annealer paddle is placed in the ‘closed’ position to prevent the gas flow from reaching the sample and heating it unnecessarily during experimental set up (**Figure 1B, left**; i.e., crystal mounting and centering, closing the experimental hutch, entering the experimental parameters into control software). After the sample is mounted, the annealer paddle is moved to the ‘open’ position (**Figure 1B, right**) via the beamline control software BluIce (McPhillips *et al*., 2002) and data collection is initiated after a short temperature equilibration delay. Control kinetic measurements showed that a J thermocouple moved from room temperature (~293 K) to a 363 K N2 stream (the highest temperature used in this work) was 360.5 K, within 5% of the desired temperature, in less than 10 seconds (not shown). We used this equilibration time prior to data collection (see below).

**Figure 1.**
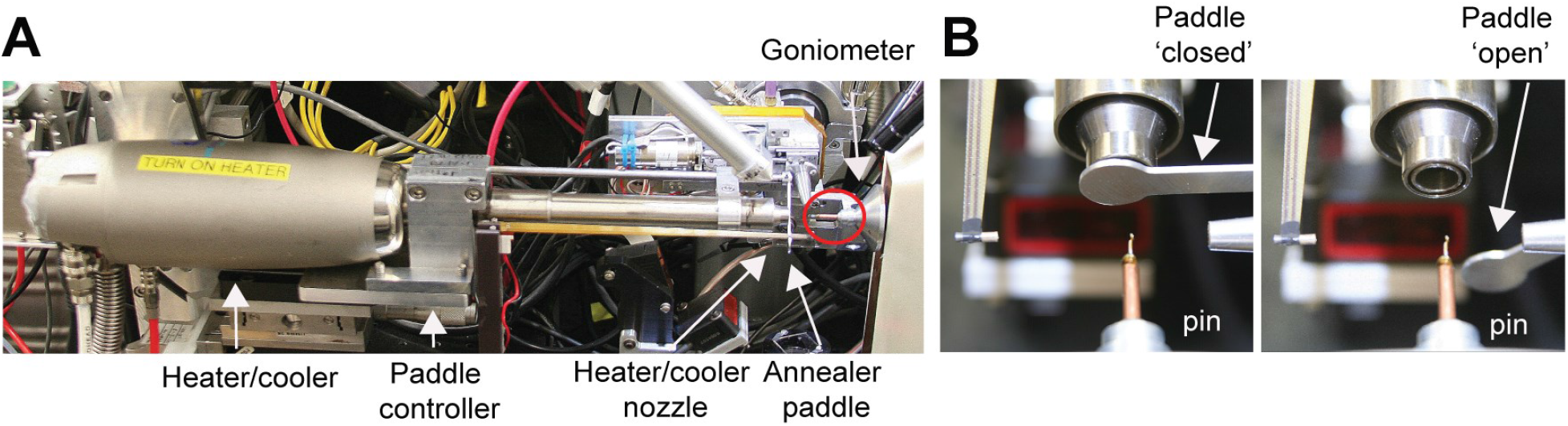
Experimental setup for X-ray diffraction data collection at and above room temperature. **A.** The Oxford Cryosystems heater/cooler mounted on the annealer device working in a ‘sample protective mode’ during preparation stages. The pin holding the sample (circled in red) is co-axially aligned with the heater/cooler nozzle. The X-ray beam and the X-ray detector are orthogonal to the sample/heater line. **B.** The annealer paddle blocks N2 in the ‘closed’ position (left) and allows the N_2_ gas to reach the crystal mounted on the pin in the ‘open’ position (right).

The diffraction data obtained provided additional independent evidence that the crystal temperature increased with increasing the N_2_ gas temperature (see **Figure 2** of Results).

**Figure 2.**
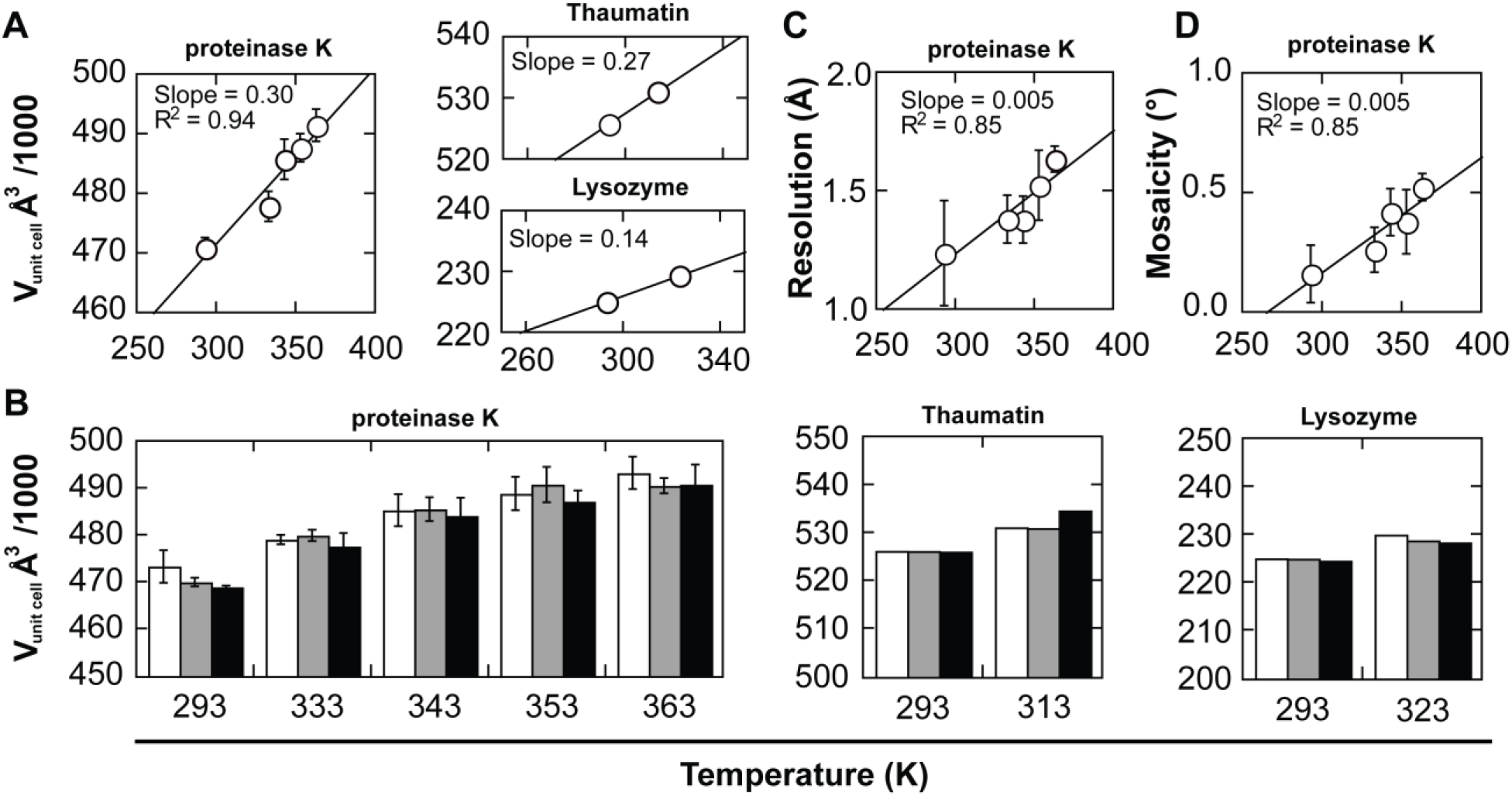
Unit cell volume, resolution, and mosaicity analysis. **A.** Proteinase K (left), thaumatin (right, top), and lysozyme (right, bottom) unit cell volume (Vunit cell) increases with increasing temperature. For proteinase K, mean Vunit cell and associated standard deviations from independent datasets obtained from three (293 K) and four (333–363 K) independent crystals at each temperature. For thaumatin and lysozyme, V_unit cell_ is obtained from a single crystal at each temperature. **B.** For proteinase K (left), thaumatin (middle), and lysozyme (right), Vunit cell does not change significantly during data collection. For proteinase K mean V_unit cell_ and associated standard deviations obtained from images 1-10 (white bars), from images 91-100 (grey bars), and from images 361-370 (black bars) (same crystal orientation as images 1-10 but after a complete 360° rotations during which the crystal was exposed to X-rays) for datasets in A. For thaumatin and lysozyme V_unit cell_ are obtained from a single crystal at each temperature. **C.** Proteinase K average dataset resolution (cut-off ~ 0.3 CC1/2) decreases with increasing temperature. **D.** Proteinase K average dataset mosaicity (images 1-10, 2.0 Å resolution cut-off) increases with increasing temperature. Mean resolutions (C) and mosaicities (D) and associated standard deviations for datasets in A. See Tables S2-S5 for values used in these plots.

### X-ray dose

At cryo temperatures the X-ray dose required to halve the diffraction intensity from an average protein crystal is about 30-40 MGy (Owen *et al*., 2006; Paithankar & Garman, 2010), and 10 MGy X-ray doses were shown to decrease diffraction resolution by 1 Å (Howells *et al*., 2009). As protein crystals are up to 300 times more sensitive to radiation damage at room temperature (Roedig *et al.*, 2016; Southworth-Davies *et al.*, 2007; Warkentin *et al.*, 2011; Warkentin & Thorne, 2010), we used doses 200-500 fold lower than the 10 MGy limit, corresponding to total doses of about 0.02-0.05 MGy for datasets of 180° total rotation. For a given total X-ray dose, higher dose rates (absorbed X-ray dose per unit of time) have been shown to extend crystal lifetimes (Southworth-Davies et al., 2007); we therefore used high dose rates of ~1 x 10^-3^ to ~4 x 10^-3^ MGy/sec (see **Table 1** and **Table S1**). While crystals suffer radiation damage even with such low doses, recent work suggested that the conformational heterogeneity in protein crystals at room temperature is not dominated by radiation damage and that specific damage does not appear before the diffraction resolution deteriorates, in contrast to observations from cryo datasets (Gotthard *et al*., 2019; Russi *et al*., 2017; Roedig *et al*., 2016).

**Table 1.**
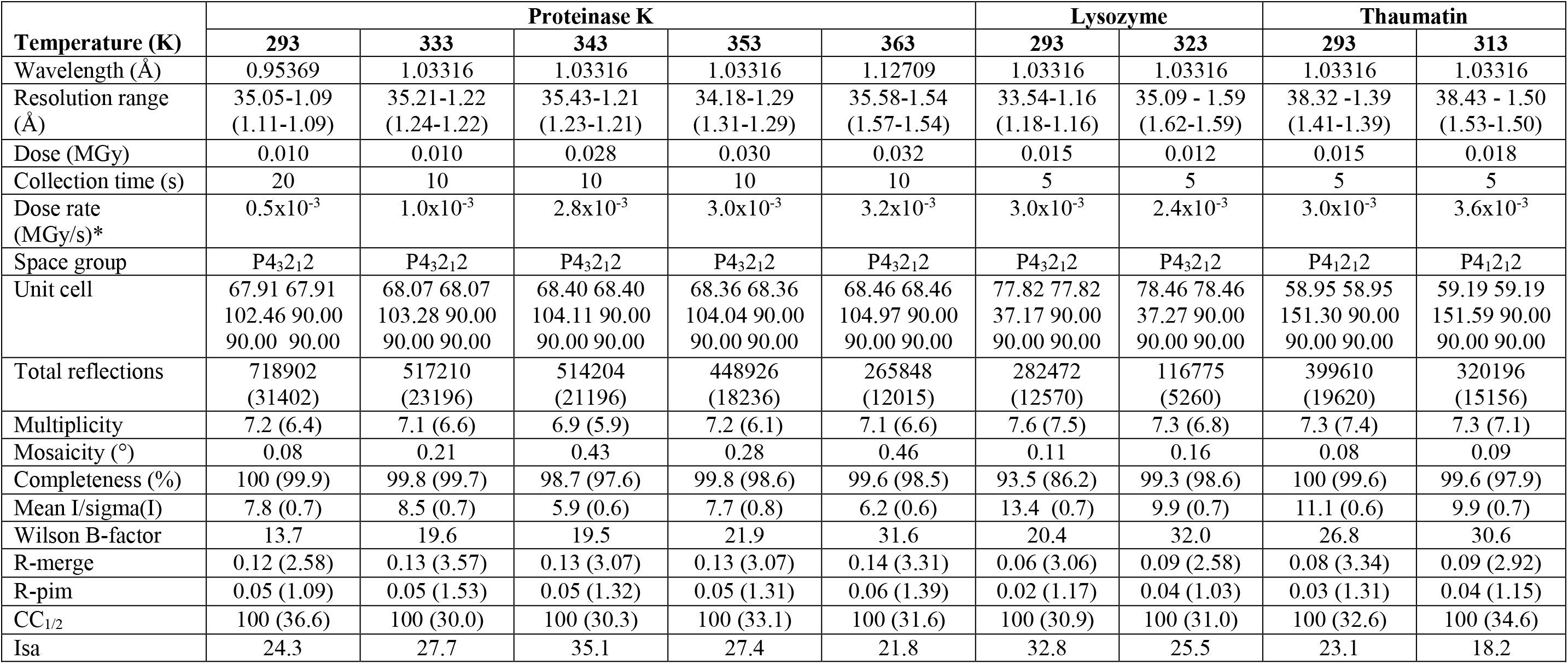
Diffraction statistics. Proteinase K, thaumatin, and lysozyme crystals diffraction statistics are reported for datasets of 100° total rotations, which were sufficient for high completeness. Values in parenthesis are for the highest resolution shells. Unit cell parameters were obtained using images from the entire 100°. All statistics were obtained from Aimless (Evans & Murshudov, 2013), with the exception of Wilson B-factor, ISa and CC1/2, which were obtained from XSCALE (Kabsch, 2010). Statistics for datasets used in Figure 2 are reported in Table S1. *Eiger 16M readout time between frames is 3 μsec (Casanas *et al.*, 2016), which corresponds to a 300 μsec total readout time for a complete dataset of 100 images, which is negligible relative to the 5-20 sec total collection time.

### Minimizing time-dependent X-ray damage at room temperature

At room temperature, the X-ray induced damage has a time component, such that damage continues even after the X-ray source has been turned off (Blundell & Johnson, 1976; Warkentin *et al*., 2011), in particular when, after collecting a few test diffraction images, the crystal is left on the goniometer while the experiment is set (usually on the order of minutes) and when the shutter is closed after collecting a frame on a charge-coupled device (CCD) detector and before it is opened again for the next frame (“readout time”, typically between a few seconds to dozens of seconds between frames).

To circumvent these limitations and reduce time-dependent X-ray damage and associated diffraction intensity decay, we eliminated the initial X-ray test exposures that are traditionally used for cryo data collection (Dauter, 1999). Instead, we implemented a fast data collection strategy in which a total rotation range of 180° can be collected immediately following thermal equilibration of the crystal. A complete dataset can be obtained for most protein crystal types and initial crystal orientations from 180° total rotation (Dauter, 1999, 2017). For the protein crystals in this work, we used 100° total rotation as this range provided complete, high-resolution datasets (**Tables 1** and **S1**). To further reduce time-dependent X-ray damage and minimize the total data collection time, we used rotation images of 1° with exposure times of 0.05–0.2 sec per image, allowing a complete dataset to be collected within ~3-36 sec, depending on the symmetry of the crystal (Dauter, 1999, 2017). The 0.05–0.2 sec (5 to 20 Hz) exposure times were enabled by the use of high-frequency frame rate photon counting detectors (the Eiger 16M detector was used in this work; the 0.05-0.2 sec exposure times enable the use of both the Eiger 16M and Pilatus 6M detectors, expanding the applicability of the approach to a larger number of beamlines), with the x-ray’s shutter closing and opening after the recording of each frame eliminated (i.e., ‘shutterless’ mode) to further reduce the total experimental time and the time-dependent X-ray damage (Brönnimann *et al*., 2003). In our experiments, the full potential of the Eiger 16M detector (133 Hz) was not achieved due to current flux limitations at BL14-1 (1.7×10^11^ photons/s at 10.5 keV). We estimate that a complete 180° dataset could be obtained in ≤1 sec if the X-ray flux were increased 10-15 fold, which could have additional benefits with respect to outrunning radiation damage (Warkentin *et al*., 2011). Careful evaluation of the hardware and software capabilities will be required if higher rotational speeds and data collection frequencies are to be routinely used for collecting high-quality data (Diederichs, 2010; Casanas *et al*., 2016).

The goal of the approach herein is to collect high-quality, complete datasets at and above room temperature. To increase the likelihood of success in collecting high-quality diffraction data for a given project, a few crystals of similar size and with similar diffraction properties are required and the crystals need to be prepared in a standardized manner for data collection (see *‘Preparing crystals for data collection’*). Slight adjustments in the experimental setup may be needed and implemented (see below), but our experience suggests that uniform work practices increase the success rate. As discussed above, collection of high-quality, complete datasets at and above room temperature requires small X-ray doses, and we applied doses on the order of 0.02-0.05 MGy. Because the total absorbed X-ray dose by a crystal during diffraction data collection directly depends on the experimental parameters (beam intensity, beam size, rotation range, and collection frequency), these parameters need to be set prior to data collection to achieve the desired dose. Because, as discussed above, fast data collection is required to outrun time-dependent radiation damage effects, exposure time per image should be short (0.05-0.2 s herein or faster at brighter beamlines) and to cover a rotation range of 180° (or less, depending on crystal symmetry), per-image rotations were set to 1°. The beam intensity to achieve a desired dose can in principle be estimated using the program RADDOSE, which would also require information about the crystal, unit cell size, and solvent and protein content, among others. (Bury *et al*., 2018; Zeldin *et al*., 2013). While such dose estimation could be used to directly establish parameters for data collection, in practice we find that using a test crystal diffraction together with initial dose estimates allows us to adjust experimental parameters as needed to collect within the 0.02-0.05

MGy range. While 2-fold X-ray dose over- or underestimation will generally not significantly impact data collection and quality (and there appears to be a 2-fold uncertainty associated with estimates in general (Holton, 2009)), severe overestimation or underestimation of the dose will lead to weak, suboptimal diffraction or excessive damage and incomplete diffraction datasets, respectively, outcomes that can be quickly detected as the diffraction data are analyzed.

### Preparing crystals for data collection

Immediately prior to data collection, the outer layer of the crystal’s aqueous mother liquor was exchanged to an inert oil (paratone-N) in the following way: the drop containing the crystals was completely covered with an excess of the paratone-N oil to prevent crystals in the drop from dehydrating (Hope, 1988). Within the drop, each crystal used for data collection was transferred from the mother liquor to the oil while the aqueous layer was stripped (Hope, 1988). This procedure eliminates the potential risk from crystal dehydration by exposing the crystal to air, as is often done when crystals are prepared for data collection using alternative devices (e.g. polymer sleeves, glass capillaries). In addition, this procedure utilizes uniformly all crystals from a drop. Due to its high viscosity and hydrophobicity, paratone-N acts as an immiscible barrier for water and significantly reduces evaporation (Hope, 1988, 1990; Pflugrath, 2015). Inside an oil drop the aqueous layer on the crystal surface was removed with a nylon loop and the oil-covered crystals were mounted on Dual-Thickness MicroLoops LD™ and MicroGrippers™ loops (Mitegen). Excessive oil was removed using a second nylon loop until only a thin coating remained, as any material in the beam would increase the background scattering. The pins were mounted on the goniometer for thermal equilibration followed by data collection (see *‘Achieving high-temperature capabilities and temperature control*’).

### Diffraction data processing

All diffraction datasets were processed using the XDS package (Kabsch, 2010) and the programs Pointless (Evans, 2006) and Aimless (Evans & Murshudov, 2013), as implemented in the *autoxds* in-house processing scripts at SSRL (https://smb.slac.stanford.edu/facilities/software/xds/).

## Results

Typically, the goal of X-ray diffraction experiments at and above room temperature is to obtain information about protein conformational heterogeneity and solvent structure for proteins whose overall structure is known (Dunlop *et al*., 2005; Fischer *et al*., 2015; Keedy *et al*., 2014, 2015; Thomaston *et al*., 2017; Woldeyes *et al*., 2014). Therefore, the main requirements for datasets obtained across physiological temperatures are high resolution and full completeness of the diffraction data, and we have developed the experimental approach herein accordingly.

To evaluate our experimental approach, we collected data from proteinase K, thaumatin, and lysozyme crystals. We obtained single-crystal X-ray diffraction datasets at and above room temperature with estimated absorbed doses of about 0.01-0.03 MGy (**Tables 1** and **S1**). All datasets were of outstanding quality, as evidenced by the very high resolutions and excellent diffraction statistics (**Tables 1** and **S1)**. The maximum temperature of data collection was limited only by the physical stability of the crystals at the desired temperature (see below).

For proteinase K, we could obtain complete high-resolution diffraction datasets up to 363 K, while the highest temperatures for thaumatin and lysozyme were 313 K and 323 K, respectively (**Tables 1** and **S1**). Above these temperatures we observed an abrupt loss of diffraction. Previous studies, in which X-ray diffraction datasets were collected at increasing temperatures, reported expansion of crystal unit cells with temperature (Keedy *et al*., 2015; Kurinov & Harrison, 1995; Tilton *et al*., 1992). To determine if we observe similar unit cell thermal expansion, we collected additional datasets for proteinase K crystals within the 293–363 K temperature range; we used several crystals at each temperature and collected an independent and complete dataset from each crystal (**Tables 1** and **S1**). **Figure 2A** shows that the average unit cell expands with temperature, consistent with previous observations, and suggesting that the desired temperature has been achieved. The observed slope of 0.3 indicated that the proteinase K unit cell expands with 300 Å^3^ K^-1^, and we observed slopes of 270 Å ^3^ K^-1^ and 140 Å ^3^ K^-1^ for thaumatin and lysozyme, respectively, (**Figure 2B)**.

In contrast to the observed volume increase with increasing temperature, excessive dehydration of protein crystals has been shown to correlate with large decreases in unit cell volume (Atakisi *et al.*, 2018). To determine if dehydration occurred during our data collection, we compared unit cell volumes (V_unit cell_) from different stages of the experiments. For the proteinase K temperature series, we compared mean volumes from independent datasets, each collected from independent crystals. To evaluate the extent of changes in V_unit cell_ during data collection, we compared V_unit cell_ calculated from images 1-10 and 91-100 from each dataset (the first and last 10° from each dataset, respectively). To evaluate if unit cell changes have occurred after collection of 360° of total rotations from each crystal and compare V_unit cell_ obtained from the same crystal orientation, we also compared the V_unit cell_ obtained from images 1-10 and 361-370. **Figures 2B** shows that V_unit cell_ from images 1-10 from each dataset (white bars) are similar to the V_unit cell_ either from images 91-100 from each dataset (grey bars) or from images 361-370 (black bars). The small variations in V_unit cell_ are consistent with the previously estimated ~0.2% uncertainties in the determination of unit cell dimensions (Dauter & Wlodawer, 2015). These observations suggest that no significant dehydration occurred during data collection.

**Figure 2C** shows decreasing resolution across temperature for proteinase K crystals, with a slope of 0.005 Å K^-1^ and still high resolution of 1.54 Å at 363 K; similar decreases in resolution are observed for thaumatin and lysozyme (**Table 1**). Because all proteinase K datasets were obtained with similar data collection parameters, from several independent crystals per temperature, and from crystals with similar size and shape, it is unlikely that the observed dependency is fortuitous and caused by random crystal-to-crystal variation.

In the simplest scenario, diffraction resolution decay could be caused by increased sensitivity to radiation damage with temperature, such that crystal diffraction decays faster at higher temperatures, and future experiments can be designed to evaluate this model. The decay in diffraction resolution could alternatively be caused by increased crystal disorder due to increased motions within the crystal with increasing temperature. If this were the case, then we would expect to see a clear trend of increasing crystal mosaicity with increasing temperature. **Figure 2D** shows such a clear trend in increasing mosaicity with temperature, with a slope of 0.005 ° K^−1^, identical to the slope of 0.005 Å K^−1^ observed in **Figure 2C**. This observation supports a direct link between mosaicity and resolution and suggests that the decrease in resolution with temperature is caused by increasing crystal disorder, as captured by mosaicity. Increased crystal disorder could originate from increasing conformational heterogeneity within the crystal unit cell. Future work will test this latter model by evaluating proteinase K motion with increasing temperature.

## Discussion

A complete quantitative and predictive understanding of biology requires an ability to predict the energetics of protein folding, ligand binding, and function. While traditional X-ray crystallography structures have been and remain invaluable in biology and medicine, they do not provide the conformational ensemble information needed to relate structure to energetics. Recent advances in room-temperature X-ray crystallography have demonstrated the ability to obtain ensemble information and relate this information to function (Dunlop *et al*., 2005; Fraser *et al*., 2009; Fraser & Jackson, 2011; Keedy *et al*., 2015), and recent technical and methodological advances in data collection indicated that high-temperature X-ray data collection is possible but further developments are required to achieve the high resolutions needed to obtain ensemble information at temperatures above room temperature (Rajendran *et al*., 2011).

Here we develop and demonstrate a robust and widely applicable method for collecting high quality X-ray diffraction data across physiological temperatures at a synchrotron beamline from single crystals. We collected high-resolution X-ray diffraction data of outstanding quality in the 293–363 K temperature range from proteinase K, thaumatin, and lysozyme crystals. This is the first time, to our knowledge, that data beyond 2.0 Å resolution have been collected above 333 K and that complete and high-resolution X-ray diffraction datasets have been collected at 363 K. Further, while the crystals used here diffracted to high resolutions (1.0–1.5 Å), it is possible to obtain meaningful biological information at lower resolutions (≤2.5 Å) where major side chain alternative rotameric states and bound water molecules are still identified in the electron density, thus significantly expanding the applicability of the approach (Wlodawer *et al*., 2008; Lang *et al*., 2010). Most impactfully, the approach presented here will allow high-quality X-ray data to be obtained more routinely at physiological temperatures.

Our method was implemented at SSRL beamline 14-1 but can be readily implemented at other SSRL beamlines and other synchrotrons. The crystal annealer device to control temperature equilibration can be built and adapted to most beamline setups in the matter of days. Fast data collection can be achieved either using the Eiger 16M detector (in this work) (Casanas *et al*., 2016), or the Pilatus 6M detector (Broennimann *et al*., 2006) available at most synchrotrons (including at SSRL beamlines 9-2 and 12-2). The X-ray flux of at BL14-1 (1.7×10^11^ photons/s at 10.5 keV) is rather standard for protein X-ray beamlines (http://biosync.rcsb.org/), and larger beams required to match larger crystals are achievable by adjusting the X-ray optic instruments.

The method described herein is complementary to room temperature serial crystallography XFEL (SFX) or synchrotron (SMX) approaches that use microcrystals, with the advantage of potentially delivering higher resolutions from single crystals and excluding potentially complicating effects from non-isomorphous multi-crystal averaging, limited XFEL facilities beam time availability, long collection and processing times, but with the disadvantage of datasets not being completely radiation damage-free. Nevertheless, recent research indicated that radiation damage does not significantly impact the conformational heterogeneity in protein crystals at room temperature (Gotthard *et al*., 2019; Russi *et al*., 2017; Roedig *et al*., 2016). In addition, data collection times can be further decreased to ≤1 sec for a complete dataset at beamlines with higher X-ray fluxes. This faster collection could outrun a significant fraction of the remaining damage (Warkentin *et al*., 2011), and shorter data collection times can allow more data to be collected at high-demand high-performance synchrotron beamlines.

The ability to robustly and efficiently collect X-ray diffraction data from single crystals at and above room temperature and obtain high-quality diffraction data also opens new opportunities for structural biologists and protein biochemists. First, it will be possible to obtain experimental phasing information directly at room temperature. While currently room temperature data is collected for proteins for which the structure has previously been solved at cryo temperatures, experimentally solving and obtaining conformational heterogeneity information for a new structure in a single experiment will reduce experimental time and modeling efforts. Similarly, the ability to obtain accurate experimental phases directly at room temperature can also help remove potential bias carried over from molecular replacement models obtained at cryo temperature by solving the structure and obtaining conformational heterogeneity information under the same conditions. Our preliminary results indicate that diffraction data is of high enough quality to allow native (SAD) phasing (manuscript in preparation), which provides additional evidence for the outstanding quality of the data obtained using this approach. Intriguingly, the ability to obtain high-quality diffraction data at high temperatures, as developed and presented in this work, may also enable the direct observation of structural and conformational heterogeneity changes on atomic level that precede unfolding events in proteins.

## Acknowledgments

Use of the Stanford Synchrotron Radiation Lightsource (SSRL), SLAC National Accelerator Laboratory, is supported by the U.S. Department of Energy, Office of Science, and Office of Basic Energy Sciences under Contract No. DE-AC02-76SF00515. The SSRL Structural Molecular Biology Program is supported by the DOE Office of Biological and Environmental Research and by the National Institute of Health (NIH), National Institute of General Medical Sciences (NIGMS, P41GM103393). The contents of this publication are solely the responsibility of the authors and do not necessarily represent the official views of NIH or NIGMS. We thank Lisa Dunn for beam time allocation and access. We also thank James Fraser, Henry van den Bedem, Tung-Chung Mou, Thomas Poulos, and members of the Herschlag lab for helpful discussions and comments on the manuscript. Part of this work was funded by a National Science Foundation (NSF) Grant (MCB-1714723) to DH. FY was supported in part by a long-term Human Frontiers Science Program postdoctoral fellowship.

## References

Atakisi, H., Moreau, D. W. & Thorne, R. E. (2018). Acta Crystallogr. Sect. Struct. Biol. 74, 264–278.

Berg, J. M., Tymoczko, J. L., Stryer, L., Berg, J. M., Tymoczko, J. L. & Stryer, L. (2002). Biochemistry W H Freeman.

Blundell, T. L. & Johnson, L. N. (1976). Protein Crystallography Academic Press.

Brändén, C.-I. & Tooze, J. (1999). Introduction to Protein Structure Taylor & Francis.

Broennimann, C., Eikenberry, E. F., Henrich, B., Horisberger, R., Huelsen, G., Pohl, E., Schmitt, B., Schulze-Briese, C., Suzuki, M., Tomizaki, T., Toyokawa, H. & Wagner, A. (2006). J. Synchrotron Radiat. 13, 120–130.

Brönnimann, C., Eikenberry, E. F., Horisberger, R., Hülsen, G., Schmitt, B., Schulze-Briese, C. & Tomizaki, T. (2003). Nucl. Instrum. Methods Phys. Res. Sect. Accel. Spectrometers Detect. Assoc. Equip. 510, 24–28.

Bury, C. S., Brooks-Bartlett, J. C., Walsh, S. P. & Garman, E. F. (2018). Protein Sci. 27, 217–228.

Casanas, A., Warshamanage, R., Finke, A. D., Panepucci, E., Olieric, V., Nöll, A., Tampé, R., Brandstetter, S., Förster, A., Mueller, M., Schulze-Briese, C., Bunk, O. & Wang, M. (2016). Acta Crystallogr. Sect. Struct. Biol. 72, 1036–1048.

Dauter (1999). Acta Crystallogr. D Biol. Crystallogr. 55, 1703–1717.

Dauter (2017). Protein Crystallography: Methods and Protocols, Vol. edited by Wlodawer, Dauter & Jaskolski, pp. 165–184. New York, NY: Springer.

Dauter, Z. & Wlodawer, A. (2015). Acta Crystallogr. D Biol. Crystallogr. 71, 2217–2226.

Diederichs, K. (2010). Acta Crystallogr. D Biol. Crystallogr. 66, 733–740.

Dunlop, K. V., Irvin, R. T. & Hazes, B. (2005). Acta Crystallogr. D Biol. Crystallogr. 61, 80–87.

Elias, M., Wieczorek, G., Rosenne, S. & Tawfik, D. S. (2014). Trends Biochem. Sci. 39, 1–7.

Evans & Murshudov (2013). Acta Crystallogr. D Biol. Crystallogr. 69, 1204–1214.

Evans, P. (2006). Acta Crystallogr. D Biol. Crystallogr. 62, 72–82.

Feller, G. (2010). J. Phys. Condens. Matter. 22, 323101.

Fersht, A. (2017). Structure and Mechanism in Protein Science: A Guide to Enzyme Catalysis and Protein Folding World Scientific.

Fields, P. A., Dong, Y., Meng, X. & Somero, G. N. (2015). J. Exp. Biol. 218, 1801–1811.

Fischer, M., Shoichet, B. K. & Fraser, J. S. (2015). ChemBioChem. 16, 1560–1564.

Fraser, J. S., van den Bedem, H., Samelson, A. J., Lang, P. T., Holton, J. M., Echols, N. & Alber, T. (2011). Proc. Natl. Acad. Sci. 108, 16247–16252.

Fraser, J. S., Clarkson, M. W., Degnan, S. C., Erion, R., Kern, D. & Alber, T. (2009). Nature. 462, 669–673.

Fraser, J. S. & Jackson, C. J. (2011). Cell. Mol. Life Sci. 68, 1829–1841.

Garman, E. F. & Owen, R. L. (2006). Acta Crystallogr. Sect. Wiley-Blackwell. 62, 32–47.

Garman, E. F. & Weik, M. (2017). J. Synchrotron Radiat. 24, 1–6.

Gotthard, G., Aumonier, S., De Sanctis, D., Leonard, G., von Stetten, D. & Royant, A. (2019). IUCrJ. 6, 665–680.

Halle, B. (2004). Proc. Natl. Acad. Sci. 101, 4793–4798.

Holton, J. M. (2009). J. Synchrotron Radiat. 16, 133–142.

Hope, H. (1988). Acta Crystallogr. Sect. B. 44, 22–26.

Hope, H. (1990). Annu. Rev. Biophys. Biophys. Chem. 19, 107–126.

Howells, M. R., Beetz, T., Chapman, H. N., Cui, C., Holton, J. M., Jacobsen, C. J., Kirz, J., Lima, E., Marchesini, S., Miao, H., Sayre, D., Shapiro, D. A., Spence, J. C. H. & Starodub, D. (2009). J. Electron Spectrosc. Relat. Phenom. 170, 4–12.

Huang, P.-S., Boyken, S. E. & Baker, D. (2016). Nature. 537, 320–327.

Jeremy Mark Berg, John L. Tymoczko, Lubert Stryer Biochemistry.

Juers, D. H. & Matthews, B. W. (2001). J. Mol. Biol. 311, 851–862.

Kabsch, W. (2010). Acta Crystallogr. D Biol. Crystallogr. 66, 125–132.

Keedy, D. A., Hill, Z. B., Biel, J. T., Kang, E., Rettenmaier, T. J., Brandão-Neto, J., Pearce, N. M., von Delft, F., Wells, J. A. & Fraser, J. S. (2018). ELife. 7, e36307.

Keedy, D. A., Kenner, L. R., Warkentin, M., Woldeyes, R. A., Hopkins, J. B., Thompson, M. C., Brewster, A. S., Van Benschoten, A. H., Baxter, E. L., Uervirojnangkoorn, M., McPhillips, S. E., Song, J., Alonso-Mori, R., Holton, J. M., Weis, W. I., Brunger, A. T., Soltis, S. M., Lemke, H., Gonzalez, A., Sauter, N. K., Cohen, A. E., van den Bedem, H., Thorne, R. E. & Fraser, J. S. (2015). ELife. 4, e07574.

Keedy, D. A., van den Bedem, H., Sivak, D. A., Petsko, G. A., Ringe, D., Wilson, M. A. & Fraser, J. S. (2014). Structure. 22, 899–910.

Kuhlman, B. & Bradley, P. (2019). Nat. Rev. Mol. Cell Biol. 20, 681–697.

Kurinov, I. V. & Harrison, R. W. (1995). Acta Crystallogr. Sect. D. 51, 98–109.

Marks, D. S., Hopf, T. A. & Sander, C. (2012). Nat. Biotechnol. 30, 1072–1080.

McPhillips, T. M., McPhillips, S. E., Chiu, H.-J., Cohen, A. E., Deacon, A. M., Ellis, P. J., Garman, E., Gonzalez, A., Sauter, N. K., Phizackerley, R. P., Soltis, S. M. & Kuhn, P. (2002). J. Synchrotron Radiat. 9, 401–406.

Nave, C. & Garman, E. (2005). J. Synchrotron Radiat. 12, 257–260.

Orengo, C. A. & Thornton, J. M. (2005). Annu. Rev. Biochem. 74, 867–900.

Owen, R. L., Rudiño-Piñera, E. & Garman, E. F. (2006). Proc. Natl. Acad. Sci. 103, 4912–4917.

Paithankar, K. S. & Garman, E. F. (2010). Acta Crystallogr. D Biol. Crystallogr. 66, 381–388.

Pflugrath, J. (2015). Acta Crystallogr. Sect. F. 71, 622–642.

Rajendran, C., Dworkowski, F. S. N., Wang, M. & Schulze-Briese, C. (2011). J. Synchrotron Radiat. 18, 318–328.

Roedig, P., Duman, R., Sanchez-Weatherby, J., Vartiainen, I., Burkhardt, A., Warmer, M., David, C., Wagner, A. & Meents, A. (2016). J. Appl. Crystallogr. 49, 968–975.

Russi, S., González, A., Kenner, L. R., Keedy, D. A., Fraser, J. S. & van den Bedem, H. (2017). J. Synchrotron Radiat. 24, 73–82.

Sandalova, T., Schneider, G., Käck, H. & Lindqvist, Y. (1999). Acta Crystallogr. D Biol. Crystallogr. 55, 610–624.

Siddiqui, K. S. & Cavicchioli, R. (2006). Annu. Rev. Biochem. 75, 403–433.

Southworth-Davies, R. J., Medina, M. A., Carmichael, I. & Garman, E. F. (2007). Structure. 15, 1531–1541.

Thomaston, J. L., Woldeyes, R. A., Nakane, T., Yamashita, A., Tanaka, T., Koiwai, K., Brewster, A. S., Barad, B. A., Chen, Y., Lemmin, T., Uervirojnangkoorn, M., Arima, T., Kobayashi, J., Masuda, T., Suzuki, M., Sugahara, M., Sauter, N. K., Tanaka, R., Nureki, O., Tono, K., Joti, Y., Nango, E., Iwata, S., Yumoto, F., Fraser, J. S. & DeGrado, W. F. (2017). Proc. Natl. Acad. Sci. U. S. A. 114, 13357–13362.

Tilton, R. F., Dewan, J. C. & Petsko, G. A. (1992). Biochemistry. 31, 2469–2481.

Ufimtsev, I. S. & Levitt, M. (2019). Proc. Natl. Acad. Sci. 116, 10813–10818.

Warkentin, M., Badeau, R., Hopkins, J. & Thorne, R. (2011). Acta Crystallogr. Sect. D. 67, 792–803.

Warkentin, M. & Thorne, R. (2010). Acta Crystallogr. Sect. D. 66, 1092–1100.

Wlodawer, A., Minor, W., Dauter, Z. & Jaskolski, M. (2008). FEBS J. 275, 1–21.

Woldeyes, R. A., Sivak, D. A. & Fraser, J. S. (2014). Curr. Opin. Struct. Biol. 28, 56–62.

Worth, C. L., Gong, S. & Blundell, T. L. (2009). Nat. Rev. Mol. Cell Biol. 10, 709–720.

Zeldin, O. B., Gerstel, M. & Garman, E. F. (2013). J. Appl. Crystallogr. 46, 1225–1230.

