## Supplementary file for "A Robust Method for Collecting X-ray Diffraction Data from Protein Crystals across Physiological Temperatures"

| Temperature (K) | Proteinase K | | | | | | | |
|---|---|---|---|---|---|---|---|---|
| | **293** | **293** | **333** | **333** | **333** | **343** | **343** | **343** |
| Wavelength | 0.95369 | 0.95369 | 0.95369 | 0.95369 | 0.95369 | 1.03316 | 1.03316 | 1.03316 |
| Resolution range (Å) | 34.94-1.55 (1.58-1.55) | 34.99-1.08 (1.10-1.08) | 35.28-1.50 (1.53-1.50) | 35.11-1.40 (1.43-1.40) | 35.17-1.40 (1.42-1.40) | 35.53-1.45 (1.47-1.45) | 35.29-1.40 (1.42-1.40) | 35.34-1.45 (1.47-1.45) |
| Dose (MGy) | 0.005 | 0.010 | 0.016 | 0.012 | 0.009 | 0.023 | 0.013 | 0.014 |
| Collection time (s) | 20 | 20 | 10 | 10 | 10 | 10 | 5 | 5 |
| Dose rate (MGy/s)* | $0.25x10^{-3}$ | $0.5x10^{-3}$ | $1.6x10^{-3}$ | $1.2x10^{-3}$ | $0.9x10^{-3}$ | $2.3x10^{-3}$ | $2.6x10^{-3}$ | $2.8x10^{-3}$ |
| Space group | $P4_32_12$ | $P4_32_12$ | $P4_32_12$ | $P4_32_12$ | $P4_32_12$ | $P4_32_12$ | $P4_32_12$ | $P4_32_12$ |
| Unit cell | 67.74<br>67.74<br>102.13<br>90.00<br>90.00<br>90.00 | 67.90<br>67.90<br>102.18<br>90.00<br>90.00<br>90.00 | 68.18<br>68.18<br>103.55<br>90.00<br>90.00<br>90.00 | 67.89<br>67.89<br>102.93<br>90.00<br>90.00<br>90.00 | 68.02<br>68.02<br>103.08<br>90.00<br>90.00<br>90.00 | 68.49<br>68.49<br>104.57<br>90.00<br>90.00<br>90.00 | 68.23<br>68.23<br>103.54<br>90.00<br>90.00<br>90.00 | 68.26<br>68.26<br>103.75<br>90.00<br>90.00<br>90.00 |
| Total reflections | 258136 (10602) | 731044 (31801) | 287239 (12547) | 348701 (17185) | 335008 14595 | 313164 (13391) | 357529 (17185) | 314183 (13858) |
| Multiplicity | 7.3 (6.4) | 7.1 (6.3) | 7.3 (6.5) | 7.2 (7.0) | 7.1 (6.1) | 7.0 (6.3) | 7.4 (7.2) | 7.1 (6.5) |
| Mosaicity (°) | 0.33 | 0.07 | 0.41 | 0.35 | 0.48 | 0.53 | 0.29 | 0.47 |
| Completeness (%) | 99.9 (97.8) | 100 (99.9) | 99.4 (98.6) | 99.9 (98.2) | 98.3 (97.8) | 99.7 (97.2) | 99.4 (96.8) | 99.7 (98.2) |
| Mean I/sigma(I) | 8.6 (0.7) | 8.1 (0.7) | 7.5 (0.6) | 6.0 (0.5) | 5.8 (0.7) | 5.7 (0.7) | 9.3 (0.7) | 6.0 (0.6) |
| Wilson B-factor | 25.8 | 13.7 | 25.9 | 25.1 | 23.6 | 26.3 | 24.2 | 25.0 |
| R-merge | 0.143 (2.300) | 0.115 (2.568) | 0.159 (3.482) | 0.201 (5.457) | 0.144 (2.347) | 0.139 (2.806) | 0.117 (3.415) | 0.147 (3.334) |
| R-pim | 0.056 (0.970) | 0.046 (1.085) | 0.062 (1.410) | 0.078 (2.104) | 0.054 (0.984) | 0.053 (1.184) | 0.044 (1.286) | 0.060 (1.362) |
| $CC_{1/2}$ | 99.9 (32.4) | 99.9 (33.5) | 99.9 (32.9) | 99.9 (33.3) | 99.9 (32.5) | 99.9 (34.0) | 99.9 (30.5) | 99.9 (34.1) |
| Isa | 37.9 | 21.4 | 32.6 | 16.4 | 17.0 | 16.1 | 31.2 | 25.2 |

**Table S1**. **Diffraction statistics.** Proteinase K crystals diffraction statistics are reported for datasets of 100° total rotations, as these were sufficiently complete. Values in parenthesis are for the highest resolution shells. Unit cell parameters were obtained using images from the entire 100°. All statistics were obtained from Aimless (Evans & Murshudov, 2013), with the exception of Wilson B-factor, ISa and $CC_{1/2}$, which were obtained from XSCALE (Kabsch, 2010). *Eiger 16M readout time between frames is 3 µsec (Casanas et al., 2016), which corresponds to a 300 µsec total readout time for a complete dataset of 100 images, which is negligible relative to the 10 sec total collection time.

| Temperature (K) | Proteinase K | | | | | |
|---|---|---|---|---|---|---|
| | **353** | **353** | **353** | **363** | **363** | **363** |
| Wavelength (Å) | 1.03316 | 1.03316 | 1.03316 | 1.03316 | 1.12709 | 1.12709 |
| Resolution range (Å) | 35.51-1.65 (1.68-1.65) | 35.54-1.50 (1.53-1.50) | 35.38-1.65 (1.68-1.65) | 35.61-1.65 (1.68-1.65) | 35.46-1.68 (1.71-1.68) | 35.64-1.66 (1.69-1.66) |
| Dose (MGy) | 0.014 | 0.015 | 0.015 | 0.029 | 0.032 | 0.032 |
| Collection time (s) | 5 | 5 | 5 | 10 | 10 | 10 |
| Dose rate (MGy/s)* | $2.8 \times 10^{-3}$ | $3.0 \times 10^{-3}$ | $3.0 \times 10^{-3}$ | $2.9 \times 10^{-3}$ | $3.2 \times 10^{-3}$ | $3.2 \times 10^{-3}$ |
| Space group | $P4_32_12$ | $P4_32_12$ | $P4_32_12$ | $P4_32_12$ | $P4_32_12$ | $P4_32_12$ |
| Unit cell | 68.32<br>68.32<br>104.76<br>90.00<br>90.00<br>90.00 | 68.48<br>68.48<br>104.65<br>90.00<br>90.00<br>90.00 | 68.23<br>68.23<br>104.11<br>90.00<br>90.00<br>90.00 | 68.50<br>68.50<br>105.04<br>90.00<br>90.00<br>90.00 | 68.23<br>68.23<br>104.58<br>90.00<br>90.00<br>90.00 | 68.47<br>68.47<br>105.32<br>90.00<br>90.00<br>90.00 |
| Total reflections | 218638 (9069) | 291337 (13023_ | 220099 (9752) | 216616 (8631) | 206719 (9444) | 216105 (9487) |
| Multiplicity | 7.1 (6.2) | 7.2 (6.7) | 7.5 (6.6) | 7.0 (5.9) | 7.2 (6.6) | 7.1 (6.7) |
| Mosaicity (°) | 0.51 | 0.51 | 0.42 | 0.56 | 0.50 | 0.52 |
| Completeness (%) | 99.9 (98.2) | 99.6 (98.1) | 97.7 (97.2) | 99.8 (97.3) | 99.8 (97.5) | 99.8 (96.7) |
| Mean I/sigma(I) | 6.1 (0.6) | 6.0 (0.5) | 5.8 (0.6) | 5.5 (0.4) | 6.5 (0.4) | 6.7 (0.6) |
| Wilson B-factor | 31.3 | 27.3 | 31.1 | 33.8 | 35.0 | 34.1 |
| R-merge | 0.152 (3.003) | 0.161 (4.339) | 0.174 (2.298) | 0.168 (4.714) | 0.182 (5.599) | 0.151 (4.087) |
| R-pim | 0.060 (1.271) | 0.063 (1.707) | 0.065 (0.928) | 0.067 (2.058) | 0.070 (2.162) | 0.060 (1.683) |
| $CC_{1/2}$ | 99.8 (30.1) | 99.9 (32.0) | 99.9 (36.0) | 99.8 (31.5) | 99.8 (33.1) | 99.9 (36.8) |
| Isa | 23.1 | 27.2 | 19.6 | 15.7 | 18.7 | 20.4 |

| Temperature | Average Unit cell volume (Å$^3$) | Standard deviation (Å$^3$) |
|---|---|---|
| **Proteinase K** | | |
| 363 | 491.4 | 2.7 |
| 353 | 487.7 | 2.4 |
| 343 | 485.8 | 3.3 |
| 333 | 477.8 | 2.5 |
| 293 | 470.8 | 1.6 |
| **Thaumatin** | | |
| 313 | 531.1 | N/A |
| 293 | 525.8 | N/A |
| **Lysozyme** | | |
| 323 | 229.4 | N/A |
| 293 | 225.1 | N/A |

**Table S2**. Data used for Figure 2A. The average volumes and standard deviations have been obtained from the data in Tables 1 and S1.

| | Unit cell volume (Å$^3$) | | | | | | | | |
|---|---|---|---|---|---|---|---|---|---|
| **Temperature (K)** | **Imgs 1-10** | **Ave** | **S.D.** | **Imgs 91-100** | **Ave** | **S.D.** | **Imgs 361-460** | **Ave** | **S.D.** |
| **Proteinase K** | | | | | | | | | |
| 363 | 489.78 | 493.16 | 3.45 | 488.83 | 490.43 | 1.63 | 484.09 | 490.66 | 4.26 |
| | 489.95 | | | 490.06 | | | 489.98 | | |
| | 494.98 | | | 493.14 | | | 493.15 | | |
| | 497.91 | | | 489.69 | | | 495.44 | | |
| 353 | 492.15 | 488.76 | 3.56 | 490.13 | 490.68 | 3.80 | 488.39 | 487.03 | 2.36 |
| | 491.04 | | | 489.63 | | | 489.34 | | |
| | 488.89 | | | 496.72 | | | 487.24 | | |
| | 482.94 | | | 486.22 | | | 483.16 | | |
| 343 | 486.33 | 485.21 | 3.43 | 485.91 | 485.41 | 2.56 | 482.42 | 484.02 | 3.85 |
| | 481.91 | | | 483.46 | | | 480.83 | | |
| | 490.33 | | | 489.39 | | | 490.61 | | |
| | 482.25 | | | 482.90 | | | 482.23 | | |
| 333 | 480.33 | 478.92 | 1.00 | 478.09 | 479.84 | 1.19 | 475.28 | 477.48 | 2.84 |
| | 479.30 | | | 480.19 | | | 479.39 | | |
| | 477.67 | | | 481.40 | | | 481.07 | | |
| | 478.37 | | | 479.67 | | | 474.17 | | |
| 293 | 475.12 | 473.21 | 3.48 | 469.15 | 469.88 | 0.93 | 468.55 | 468.75 | 0.35 |
| | 476.19 | | | 471.18 | | | 469.25 | | |
| | 468.32 | | | 469.29 | | | 468.46 | | |
| **Thaumatin** | | | | | | | | | |
| 293 | 526.21 | | | 526.17 | | | 526.07 | | |
| 313 | 531.13 | | | 530.98 | | | 536.67 | | |
| **Lysozyme** | | | | | | | | | |
| 293 | 224.98 | | | 224.92 | | | 224.51 | | |
| 323 | 229.98 | | | 228.76 | | | 228.33 | | |

**Table S3**. Data used for Figure 2B. Unit cell volumes have been obtained using images 1-10 (beginning of the dataset), 91-100 (end of the dataset), and 361-370 (same orientation as for images 1-10 but after a complete 360° rotation). The additional comparison with images 361-370 was used as differences in unit cell parameters between images 1-10 and 91-100 could also originate from using different crystal orientations to estimate unit cell volumes. Unit cell constants have been obtained using a 2.0-Å resolution cut-off.

| Temperature (K) | Average resolution (Å) | Standard deviation (Å) |
|---|---|---|
| **Proteinase K** | | |
| 363 | 1.63 | 0.05 |
| 353 | 1.52 | 0.15 |
| 343 | 1.38 | 0.01 |
| 333 | 1.38 | 0.10 |
| 293 | 1.24 | 0.22 |

**Table S4**. Data used for Figure 2C. The average resolutions and standard deviations have been obtained from the data in Tables 1 and S1.

| Temperature (K) | Mosaicity (°) | | |
|---|---|---|---|
| | Imgs 1-10 | Average | S.D. |
| 363 | 0.49 | 0.52 | 0.06 |
| | 0.48 | | |
| | 0.5 | | |
| | 0.62 | | |
| 353 | 0.15 | 0.38 | 0.13 |
| | 0.48 | | |
| | 0.47 | | |
| | 0.41 | | |
| 343 | 0.34 | 0.42 | 0.10 |
| | 0.51 | | |
| | 0.52 | | |
| | 0.3 | | |
| 333 | 0.35 | 0.26 | 0.09 |
| | 0.23 | | |
| | 0.12 | | |
| | 0.34 | | |
| 293 | 0.08 | 0.16 | 0.12 |
| | 0.07 | | |
| | 0.33 | | |

**Table S5.** Data used for Figure 2D. The average mosaicity and standard deviations have been obtained from images 1-10 of the diffraction data in Tables 1 and S1.
